# Auranofin resistance in *Toxoplasma gondii* decreases the accumulation of reactive oxygen species but does not target parasite thioredoxin reductase

**DOI:** 10.1101/2020.04.23.058842

**Authors:** Christopher I. Ma, James A. Tirtorahardjo, Sharon Jan, Sakura S. Schweizer, Shawn A. C. Rosario, Yanmiao Du, Jerry J. Zhang, Naomi S. Morrissette, Rosa M. Andrade

**Affiliations:** Department of Medicine, Division of Infectious Diseases, University of California Irvine; Department of Microbiology and Molecular Genetics, University of California Irvine; Department of Molecular Biology and Biochemistry, University of California Irvine

**Keywords:** gold, repurposing, anti-parasitic, resistance, redox, *Toxoplasma*, auranofin, superoxide

## Abstract

Auranofin, a reprofiled FDA-approved drug originally designed to treat rheumatoid arthritis, has emerged as a promising anti-parasitic drug. It induces the accumulation of reactive oxygen species (ROS) in parasites, including *Toxoplasma gondii*. We generated auranofin resistant *T. gondii* lines through chemical mutagenesis in order to identify the molecular target of this drug. Resistant clones were confirmed with a competition assay using wild-type *T. gondii* expressing yellow fluorescence protein (YFP) as a reference strain. The predicted auranofin target, thioredoxin reductase, was not mutated in any of our resistant *lines*. Subsequent whole genomic sequencing analysis (WGS) did not reveal a consensus resistance locus, although many have point mutations in genes encoding redox-relevant proteins such as superoxide dismutase (TgSOD2) and ribonucleotide reductase. We investigated the SOD2 L201P mutation and found that it was not sufficient to confer resistance when introduced into wild-type parasites. Resistant clones accumulated less ROS than their wild type counterparts. Our results demonstrate that resistance to auranofin in *T. gondii* enhances its ability to abate oxidative stress through diverse mechanisms. This evidence supports a hypothesized mechanism of auranofin anti-parasitic activity as disruption of redox homeostasis.

## INTRODUCTION

For more than 50 years, the mainstay of treatment for acute toxoplasmosis has been a combination of a dihydrofolate reductase inhibitor and a sulfa antimicrobial. While this is currently a critical therapeutic strategy, currently approved drugs cannot eliminate latent parasites in cysts that are found in chronically infected individuals. Moreover, these drugs have significant bone marrow toxicity, are suspected teratogens and the emergence of resistance remains a potential threat to treatment. Repurposing FDA-approved drugs will accelerate drug discovery for neglected parasitic diseases. Most recently, auranofin, a reprofiled drug that is FDA-approved for treatment of rheumatoid arthritis, has emerged as a promising anti-parasitic agent. It has antiproliferative activity against *Plasmodium falciparum* (1), *Schistosoma mansoni* (2)*, Leishmania infantum* (3) and *Entamoeba histolytica* (4) as well as other parasites with public health significance (5–11). Despite this promising broad spectrum of anti-parasitic activity, the molecular target of auranofin has only been indirectly implicated as thioredoxin reductase.

Thioredoxin reductase enzymes are found in archaea, bacteria and diverse eukaryotes and play a key role in reduction of disulfide bonds that is essential for cell replication and survival. Inhibition of human thioredoxin reductase 1 induces expression of heme oxygenase-1 to repress inflammation(12). We previously demonstrated that auranofin reduces *T. gondii* infection of host cells *in vitro* and a single dose of auranofin allows survival of chicken embryos infected with this parasite (13). It is hypothesized that the anti-parasitic activity of auranofin comes from inhibition of the thioredoxin reductase enzyme of these parasites, akin to its action on human cells (4, 8, 14). This postulated mechanism of action explains why parasites treated with auranofin accumulate ROS (4). Intriguingly, auranofin may improve infection outcome by decreasing pathogenic host inflammatory damage in addition to its direct inhibition of parasite replication.

In this paper we describe our work to investigate resistance to auranofin in *T. gondii* parasites. This approach illuminates mechanisms of resistance that might arise during its use as an anti-parasitic. In some circumstances, identification of resistance mechanisms validates proposed drug targets (*i.e*.(15)). In this study, we found that resistance was not associated with changes to the *Toxoplasma* thioredoxin reductase gene and no other single locus emerged as a consistent site underlying resistance. However, we observed that resistant clones accumulated less ROS than their wild type counterparts, demonstrating that auranofin resistance enhances oxidative stress responses.

## MATERIAL AND METHODS

### 1. Host cell and parasite cultures

Parasite clones and human foreskin fibroblasts (HFF) were grown and maintained in Dulbecco’s modified Eagle’s medium supplemented with 10% fetal bovine serum (Hyclone), penicillin and streptomycin (50 μg/ml each) and 200 μM L-glutamine. We will refer to this complete medium as D10. T. gondii were maintained by serial passage in HFF monolayers at 37°C in a humid 5% CO2 atmosphere, including RH parasites (National Institutes of Health AIDS Reference and reagent Repository, Bethesda, MD), RH tachyzoites expressing cytoplasmic yellow fluorescent protein (YFP) (kindly provided by M.J. Gubbels, Boston College, Boston, Massachusetts) (16) and Ku80 parasites knock-out parasites (ΔKu80, kindly provided by V. Carruthers, University of Michigan Medical School Dexter, Michigan) (17, 18).

### 2. Generation of *T. gondii* clones resistant to auranofin

We mutagenized *T. gondii* with ENU as previously described (19, 20). Briefly, approximately 1.5×10^7^ intracellular tachyzoites of wild-type RH strain *T. gondii* were mutagenized with ENU (N-Ethyl-N-nitrosourea, N3385-1G, Sigma Aldrich; 100-200 μg/ml) in DMEM without FBS for 2 hours at 37°C. These parasites were washed and transferred to a new confluent HFF monolayer for selection with auranofin (2.5 μM-3 μM) until normal pace of replication was observed (~2 weeks). Auranofin resistant clones were single cell cloned by limiting dilution in 2 μM auranofin. Isolated clones were propagated in D10 media with 1 μM auranofin to maintain selection pressure without cytotoxic effects on host cells (*data not shown*).

We selected ten representative *T. gondii* clones from independent ENU-generated pools to investigate resistance. To assess auranofin resistance, we modified a tachyzoite growth competition assay (21). Using wild-type YFP-expressing *T. gondii* parasites as a control, confluent monolayers of HFF cells were inoculated with approximately equal numbers (2×10^6^; 1:1 ratio) of ENU-mutagenized parasites and YFP-expressing wild-type RH parasites. Upon lysis, a new T25 flask with confluent HFF was inoculated with 0.25 ml of lysed parasites. Another 0.25 ml of lysed parasites was used for flow cytometry. The YFP fluorescence allowed us to differentiate wild-type and Aur^R^ parasites in the mixed populations to track their growth over serial passages. Host cell debris was removed by filtering of parasites with a 3 μm filter membrane. After parasites were collected by centrifugation (400g for 10 min), the pellet was resuspended in 1 ml PBS. The total number of parasites and the number of parasites expressing YFP were quantified by flow cytometry using a FACSCalibur flow cytometer (Becton Dickinson) and analyzed with Flowjo software (Tree Star). The result of three independent experiments analyzing the replication fitness advantage between each auranofin resistant clone and the wild-type parasites was analyzed with parametric paired t-tests. A two-tailed *p*-value <0.05 was considered statistically significant.

### 3. Identification of mutations in auranofin resistant *T. gondii* clones

We extracted genomic DNA (gDNA) from auranofin resistant parasites lines using the DNeasy Blood and Tissue kit (Qiagen). The Thioredoxin reductase gene (TGGT1_309730, ToxoDB.org) was amplified from gDNA samples as two PCR fragments ~3.7 kb each. The 5’ half of the gene was amplified with primers GGACCTATGAGTTGCCTCTCTG and GTCAAATCCCAATTCGCGGAG, and the 3’ half with GTCTGCTCTCTCTGTTTCGCAAG and CCTCCACATCCTCCAGAAGC. Sequencing primers are 5TrxR 1R, 5TrxR 1F and 5TrxR 3F for the 5’ half, and TrxR F 8, 3TrxR 2F, 3TrxR 3F, and 3TrxR 4F for the 3’ half (Table S1).

To determine the presence of other SNVs in the coding region of our *T. gondii* auranofin resistant clones, gDNA from 10 resistant clones and wild-type RH parasites was submitted for library construction and whole genome sequencing at the Genomic High Throughput Facility at University of California, Irvine. Briefly, genomic DNA (gDNA) was initially quantified with a Qubit DNA High Sensitivity kit. One hundred nanograms of gDNA was sheared to <300-500> bp in size using a Covaris S220 ultrasonicator (shearing conditions: duty Cycle 10%, Cycles/burst 200 and treatment time 55 sec) (Covaris, MA). The size and the quality of the resulting DNA fragments were analyzed with an Agilent DNA High Sensitivity chip. Ten nanograms of each sample’ s fragmented DNA was end-repaired, poly adenylated and ligated to a 6bp DNA barcodes (Bioo Scientific NextFlex DNA barcodes) using a PrepX Complete ILMN DNA Library kit (Takara Bio, CA). The genomic library was built using the Apollo 324 system (Takara Bio, CA) to generate 520 bps inserts. To enrich the DNA libraries, 8 cycles of PCR amplification using a Kapa library amplification kit was used (cycling conditions: 98°C for 45 s, 8 cycles of 98°C for 15 sec, 60°C for 30 s 72° C for 30 s and then 72°C for 1 min) in an Apollo 324 system. The ultimate DNA concentration was calibrated using Qubit 2.0 dsDNA HS Assay kit and analyze with an Agilent DNA High Sensitivity chip. Before sequencing, we quantified our DNA libraries using a KAPA Library Quantification kit for Illumina platforms (Roche). Each library was normalized to 2nM and then pooled equimolar amounts in the final multiplex library that was sequenced on the Illumina HiSeq 4000.

The ten auranofin resistant *T. gondii* clones and wild-type RH *T.gondii* were sequenced with HiSeq 4000 paired end 100 reads, quality analyzed, adapter and quality trimmed before alignment with the annotated reference genome of GT1 strain from ToxoDB.org (published 7/19/2013)(22). After initial quality control with FASTQC, reads were trimmed using Illumina adapter sequences with the error probability of 0.05 and reads shorter than 20 bases were discarded. Single Nucleotide Variant (SNV) calling for the haploid genome had a coverage threshold of >5 fold. Each of the 10 mutant clones were compared to wild-type RH variant track and shared variants were filtered out. Only SNVs in the coding regions of the 10 mutant clones are reported. Only genes containing SNVs with 100 % frequency and ≥50% of read count and read coverage were considered in our final analysis. Genomic DNA from each mutant clone was used to amplify the targeted SNV region of interest. The resulting amplicons ranged between 1.7-4.5kbp in size. PCR amplicons were purified prior to sequencing via a QiaQuick PCR Purification Kit (Qiagen 28106). Amplicons were sequenced by Sanger sequencing (Genewiz), using sequencing primers targeted to the SNV region (Table S1)

### 4. Generation of modified SOD2 lines

We generated a *T. gondii* clone with a 3’ knock-in YFP tag to TgSOD2 using a ligation independent cloning approach as previously described (17). The mitochondrial distribution of our clone was confirmed by fluorescent microscopy using a Zeiss Axioskop with Axiovision camera and software. We subsequently introduced the L201P mutation into the TgSOD2-YFP line using a modified CRISPR plasmid generated using plasmid pSAG1::CAS9-U6::sgUPRT as the scaffold (23); Addgene # 54467). Our plasmid had an additional DHFR-TM marker for pyrimethamine selection derived from pLoxP-DHFR-mCherry ((24), Addgene # 70147), and the two gRNA sequences were replaced with CGGTATACCGGACAGATCCG and TAGTTTCGCCCAGTTCAAGG, selected by the Benchling software (Benchling.com).

The DNA sequence used for homology directed repair (HDR) was amplified with the primers GATAGTGTGTGAAGAGCAGC and CTGCCGTTACCAACATGG from the LIC-YFP-SOD2 plasmid generated above. PCR amplicon contained the L201P mutation (CTA->CCA) generated with Q5 Site-Directed Mutagenesis kit (New England Biolab), as with the following modifications. To screen for positive CRISPR-Cas9 clones containing L201P, two silent mutations were created to generate new and unique restriction endonuclease sites close to Leucine 201: AflI on Leucine 180 (CTC->CTT, 62 bps upstream of L201P) and XcmI on Serine 208 (TCA->TCT, 22 bps downstream of L201P). The PCR amplicon contained two additional deletions corresponding to the gRNA sites on endogenous SOD2 gene, in order to avoid SpCas9 protein cleaving the amplicon for HDR. The sequence agagacccggcgg (including the PAM sequence cgg) was deleted on the HDR sequence corresponding to gRNA #1, and sequence cacttaacagagggtc (including the PAM sequence agg) was deleted on the HDR sequence corresponding to gRNA #2.

The L201P point mutation was introduced into the TgSOD2.YFP line through CRISPR-Cas9 (Figure S1). Transfection of the CRISPR-CAS9 plasmid and the HDR DNA was carried out as published previously, with minor modifications (23). Briefly, 10^7^ of freshly lysed parasites with SOD2-YFP were filtered and spun down at 400xg and 18°C for 10 min. The pellet was resuspended with Cytomix buffer (10 mM KPO_4_, 120 mM KCl, 5 mM MgCl_2_, 25 mM HEPES, 2 mM EDTA, 2μM ATP and 5μM glutathione) up to 800μl, including 7.5μg of CRISPR plasmid and 1.5μg of HDR DNA. The mixture was transfected using the ECM 630 Electro Cell Manipulator (BTX). Parasites were incubated into T25 flasks after electroporation and switched to 2 μM pyrimethamine selection 24 hours later. Transfected pools remained in pyrimethamine selection for 3 days, and then switched to D10 media without selection for one more day before single cell cloning. The gDNA for isolated clones were collected by the DNeasy Blood and Tissue kit (Qiagen) and used to PCR amplify SOD2 with either primers SOD2 F/AmpR (for SOD2-YFP) or primers SOD E2/SOD R (for SOD2 only, Table S1). PCR amplicons were digested with XcmI for screening and positive clones were sequence verified.

### 5. Measurement of ROS accumulation in *T. gondii*

To expose our parasites to oxidative stress, we harvested freshly lysed parasites and treated 5×10^4^ parasites with 500 μM hydrogen peroxide (H_2_O_2_) for 30 min at 37°C, 5% CO_2_, with or without 1 μM auranofin After incubation, ROS was detected using H_2_DCDFC (Invitrogen). Each sample was incubated with 1 μM of H_2_DCDFC and examined under a Spectramax i5, (Molecular Devices California US; multimode microplate reader SoftMax Pro 7.0.2 Software) multimode microplate reader (excitation 494 nm; emission 522 nm) at 37°C for 15 minutes. For excitation, a single flash from a UV Xenon lamp was used for each well, and emission signals were recorded with a sensitivity setting of 100. The values are presented as relative fluorescence units. We compared ROS accumulation of each representative auranofin resistant *T. gondii* clone to that of wild-type parasites by an unpaired t-test in three independent experiments. A two-tailed *p*-value <0.05 was considered statistically significant.

### 6. Protein alignment and structural models

The predicted amino acid sequences for TgSOD2 (TGGT1_316330) and TgRNR (TGGT1_294640) were submitted to the SwissModel site to identify the optimal available protein structures for modeling (25–29). The *Toxoplasma* sequences were threaded onto the structure of *Plasmodium knowlesi* SOD (PDB 2AWP) (30) and human RNR (PDB 3HND) (31). Clustal Omega was used to align the amino acid sequences and EMBOSS was used to calculate percent identity and similarity. Information on MS peptide coverage was obtained from the ToxoDB database (22).

## RESULTS

### 1. Aur^R^ *T. gondii* lines have a growth advantage over wild type parasites

To verify auranofin resistance of the selected clones, we adapted a previously described competition assay (21). This was used in place of a normal dose-response assay because at auranofin concentrations higher than 3 μM, the fibroblast monolayer begins to disrupt (data not shown) making the plaque assays not feasible. To confirm resistance to auranofin in the Aur^R^ *T. gondii* lines, we assessed their growth in competition with YFP-expressing RH strain parasites in control media and in media with 2.5-3.0 μM auranofin. In competition assays, the inoculating parasite population is composed of 50% YFP-expressing wild type (sensitive) parasites and 50% of an Aur^R^ line verified by flow cytometry. The relative contribution of the individual populations was measured after each cycle of lytic growth in serial culture. If a parasite line increases to >50% of the population, it exhibits a growth advantage over the competing line. In the absence of auranofin, Aur^R^ lines grew comparably or less well than the parental RH line that does not express YFP (Figure 1A). However, in the presence of 2.5 μM auranofin, all Aur^R^ lines displayed significant growth advantage over YFP-expressing wild-type RH parasites (Figure 1B). The observations indicated that day 6 is the optimal day to capture the growth advantage of resistant lines under auranofin selection because YFP-expressing wild-type parasites were eliminated from the culture. Given these results, we measured the ability of a larger set of Aur^R^ lines to compete with wild-type YFP-expressing RH strain parasites in the absence or presence of auranofin by quantifying their contribution to the culture at 6 days (Figure 2). These results show that 9 of the 10 Aur^R^ lines outcompete the sensitive YFP-expressing RH line in the presence of 2.5 μM auranofin. Line 6 has a distinct profile: it grows slowly in both control and auranofin media, suggesting that it may evade auranofin-mediated killing by protracted replication.

**Figure 1.**
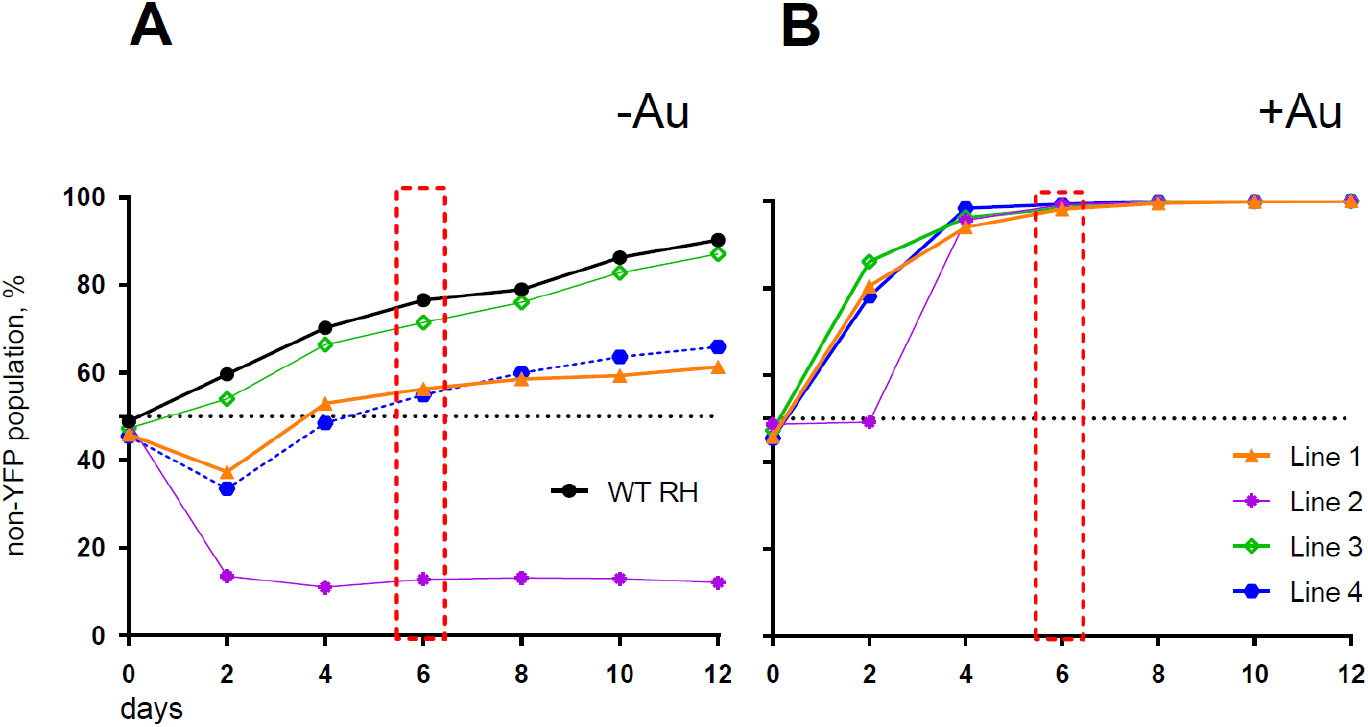
Aur^R^ lines have a growth advantage under selective pressure with auranofin. (A) In the absence of auranofin, individual resistant lines grow as well or less well than the parental RH strain. (B) In the presence of 3 μM auranofin, wild-type parasites are eliminated by day 6 (red box).

**Figure 2.**
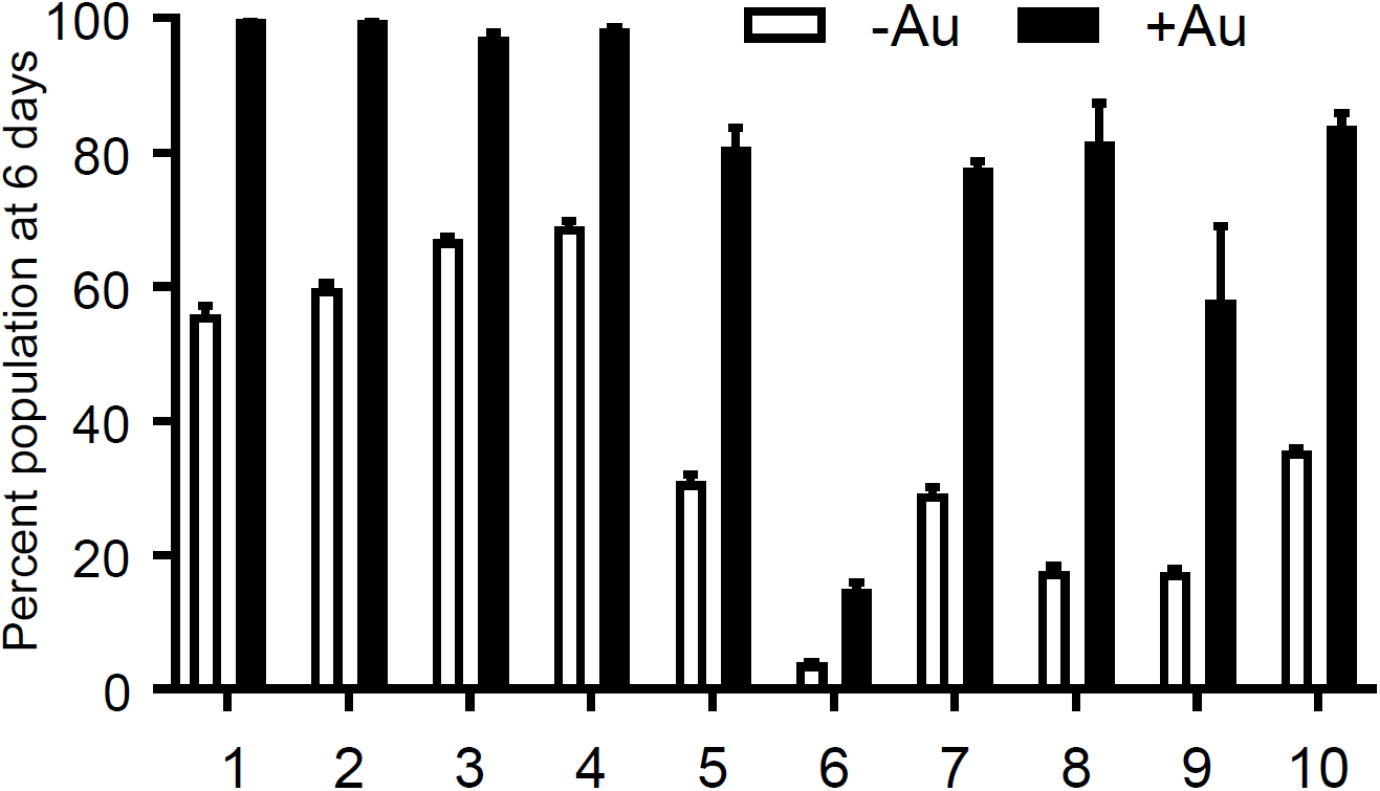
Resistant *T. gondii* lines outcompete wild type parasites under auranofin selection. Aur^R^ lines are represented as percentage of the total population of parasites after 6 days in competition with YFP-expressing wild type (sensitive) parasites, no drug media (white) and 2.5 uM auranofin (black). Mean difference between parasite population in the absence of auranofin and in the presence of auranofin: 40.6; SD 14.6; two-tailed unpaired t-test *p*<0.001.

### 2. Aur^R^ lines do not harbor mutations in the thioredoxin reductase gene

Considering that thioredoxin reductase (1–4) is the proposed target of auranofin, we hypothesized that resistance in *T. gondii* would develop through mutations to the parasite thioredoxin reductase gene (TgTrxR, TGGT1_309730). Therefore, we amplified and sequenced the TgTrxR gene in 53 lines derived from parallel (independent) selections (not shown). We did not detect mutations (SNVs) in the thioredoxin reductase gene of any of the lines, indicating that auranofin resistance does not arise through modifications to TgTrxR that directly influence auranofin binding or TgTrxR enzyme activity.

### 3. Aur^R^ lines accumulate less ROS

The anti-parasitic activity of auranofin relies on its ability to induce accumulation of ROS in parasites (4). We exposed wild type RH strain parasites and four representative Aur^R^ lines to hydrogen peroxide (H_2_O_2_) for 30 minutes before measuring ROS accumulation with dichloro-fluorescein (H_2_DCDFC). All Aur^R^ lines displayed decreased accumulation of ROS relative to the parental RH line in both the absence and presence of auranofin (Figure 3).

**Figure 3.**
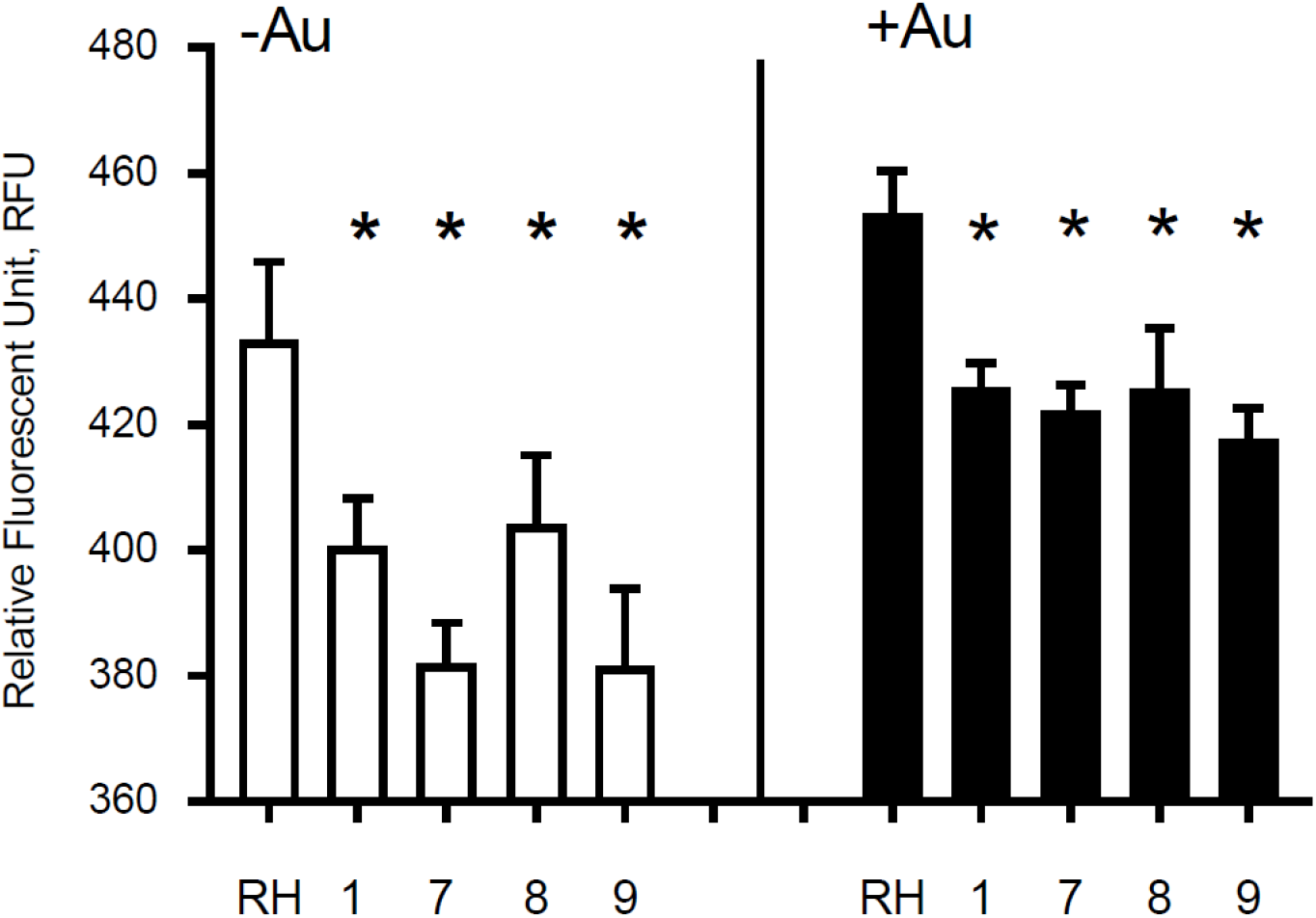
Auranofin-resistant *T. gondii* parasites accumulate less ROS. Freshly lysed wild-type parasites were treated with 500 μM hydrogen peroxide (H_2_O_2_) for 30 min and accumulation of reactive oxygen species (ROS) was assayed with dichloro-fluorescein H_2_DCDFC. Auranofin-resistant clones accumulated less ROS after H_2_O_2_ treatment compared to the wild-type RH parasites in the presence of H_2_O_2_ (white) or H_2_O_2_ + 1 μM auranofin (black). (*) *p*<0.05 when compared to RH or to RH+ Auranofin (Au).

### 4. Aur^R^ lines harbor diverse mutations that introduce point mutations

Given that we did not observe mutations in *T. gondii* thioredoxin reductase, we performed whole genome sequencing (WGS) analysis of 10 independent *T. gondii* Aur^R^ lines. Our goal was to determine whether these *T. gondii* parasites harbor a common mutation(s) that underlies resistance to auranofin. Analysis of coding sequences with 100% SNVs frequency and at least 50-fold coverage identified a variety of ~5 non-synonymous SNVs in the exons of each line. Selected SNVs observed in the WGS were validated by PCR amplification and Sanger sequencing (primers in Table S1). We chose to specifically investigate the Aur^R^ line 8 to consider which SNV was most likely to underlie auranofin resistance. Line 8 has 12 SNVs that introduce amino acid substitutions to diverse protein coding genes (Table 1). We used the community annotated ToxoDB database and BLAST analysis to investigate these loci. In particular, the availability of RNA-seq datasets permitted us to evaluate whether loci were expressed in the tachyzoite stage. In order to assess the significance of expressed loci, we used BLAST analysis to look for protein homologs and significant motifs and collected RNA-seq values and CRISPR screen scores for available loci. This information led us to focus on 2 of the loci (TGGT1_316330 and TGGT1_294640) as most likely resistance conferring. These loci encode a superoxide dismutase (SOD2) and the large subunit (R1) of ribonucleotide reductase (RNR). Both enzymes have strong fitness conferring effects in the deposited CRISPR screen data and well-conserved metabolic roles.

**Table 1.** SNVs in Aur^R^ line 8

| Table 1: SNVs in Aur<sup>R</sup> line 8 |  |  |  |  |  |  |  |
| --- | --- | --- | --- | --- | --- | --- | --- |
| Rank | Mutation | Locus | Predicted protein | CRISPR | *RNA-seq | Motif | Index |
| 1 | *L201P* | TGGT1_316330 | Superoxide dismutase SOD2 | -4.09 | 76.99 | SOD motif | 2.50 |
| 2 | *L166Q* | TGGT1_294640 | Ribonucleoside-diphosphate reductase | -4.79 | 40.9 | R1 subunit, ribonucleotide reductase | 2.29 |
| 3 | *Y95C* | TGGT1_233460 | SAG-related sequence SRS29B | -0.07 | 1809.46 | Major surface antigen p30 | 2.10 |
| 4 | *Y637H* | TGGT1_268870 | Tetrcopeptide repeat-containing | -1.66 | 38.52 | Predicted dehydrogenase | 1.81 |
| 5 | *P1357-S1358Δ* | TGGT1_310350 | PGAP1 family protein | -2.41 | 11.67 | alpha/beta-hydrolase/PGAP1-like protein | 1.45 |
| 6 | *D487G* | TGGT1_289820 | TBC domain-containing protein | -1.14 | 15.49 | Rab-GTPase-TBC domain | 1.25 |
| 7 | *L606L* | TGGT1_271290 | Hypothetical protein | -2.02 | 8.1 | Alternative splicing regulator | 1.21 |
| 8 | *F424L* | TGGT1_234250 | Hypothetical protein | -2.02 | 6.39 | SWAP/MAEBL | 1.11 |
| 9 | *A574G* | TGGT1_290910 | Hypothetical protein | -1.39 | 7.63 | No putative conserved domain | 1.03 |
| 10 | *D1983del* | TGGT1_211310 | Hypothetical protein | -0.69 | 12.86 | RCC1 repeat | 0.95 |
| 11 | *F563S* | TGGT1_260490 | Hypothetical protein | -1.62 | 5.34 | Herpes BLLF1 | 0.94 |
| 12 | *P66P* | TGGT1_225340 | Hypothetical protein | 0.73 | 24.06 | No putative conserved domain | N/A |
| *Transcriptome Gregory series at 2 hrs RH (log 2 FPKM+1). FPKM= Fragments Per Kilobase Of Exon Per Million Fragments Mapped<br>N/A: Not calculated due to dispensable (positive) CRISPR phenotype index<br>index Formula=Sum of Log10 of (-CRISPR phenotype index)+log10 (RNAseq expression) |  |  |  |  |  |  |  |

RNR removes the 2’-hydroxyl group of the ribose ring of nucleoside diphosphates to form deoxyribonucleotides from ribonucleotides which are used in DNA synthesis while SOD inactivates damaging superoxide (O_2_^-^) radicals by converting them into molecular oxygen and hydrogen peroxide. Since both TgRNR and TgSOD2 are conserved proteins, we threaded the amino acid sequence onto homologous crystal structures to locate the position of the amino acid substitutions. TgSOD2 is a Fe-type SOD and was previously demonstrated to localize to the tachyzoite mitochondrion (32, 33). SODs form dimers with shared coordination of two iron (FeIII) cofactors. The L201P mutation is located near to the key iron-coordinating residues: H111, E249 and H160 (Figure 4A) and is located in an α-helix. Substitution of a proline at this site likely causes the helix to bend, perhaps altering SOD enzymatic properties. TgRNR aligns well with human ribonucleotide reductase 1 (61.7% identity and 78.2% similarity). However, there is a 59 amino acid N-terminal extension to the *Toxoplasma* protein, with the conserved sequence beginning with M60 (Figure 4B). This leader does not represent mis-annotation of the N-terminus of the protein: there are MS peptides mapping to this sequence, especially in samples that are from monomethylarginine and phosphopeptide-enriched datasets in ToxoDB. This is particularly interesting as arginine methylation is a modification associated with proteins that have roles in transcriptional regulation, RNA metabolism and DNA repair. RNR use free-radical chemistry to convert ribonucleotides into deoxyribonucleotides. Significantly, ribose reduction requires generation of a free radical and RNR requires electrons donated from thioredoxin. Moreover, superoxide has been shown to inactivate RNR and yeast lacking either cytoplasmic or mitochondrial SODs have increased susceptibility to oxidative stress and RNR inactivation. The L166Q mutation is in a solvent exposed loop (Figure 4C), that is poorly conserved and not known to contribute to enzyme interactions.

**Figure 4.**
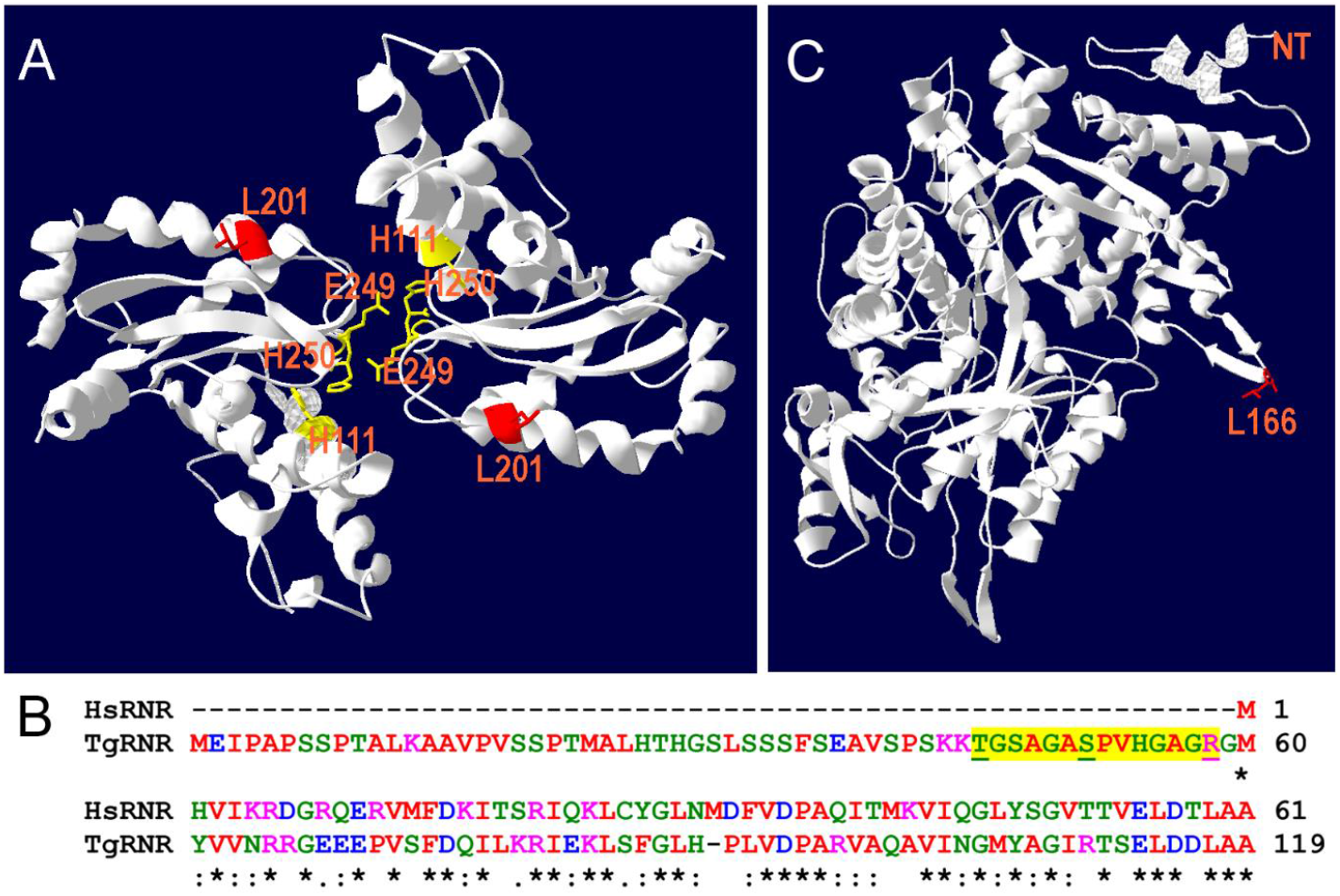
Location of substitutions in TgSOD2 and TgRNR. (A) SOD forms dimers that coordinately bind to iron cofactors at their interface. The *Toxoplasma* amino acid sequence was threaded onto a *Plasmodium knowlesi* protein structure (PDB 2AWP) to create a model for TgSOD2. The mutation at position 201 (red) is near to the end of an α-helix; substitution of a proline for leucine at this location likely causes the helix to bend. This mutation is also proximal to key iron-coordinating residues H111, E249 and H160. (B) Alignment of the *Toxoplasma* and human RNR sequences indicates that TgRNR has a novel 59 amino acid N-terminal extension. MS peptides (yellow highlight) in ToxoDB map to this sequence with S and T phosphorylated, and R monomethylated (underlined). (C) The *Toxoplasma* amino acid sequence for RNR was threaded onto the human RNR structure (PDB 3HND) to create a model for TgRNR. The L166Q mutation is in a solvent exposed loop, that is poorly conserved and not known to contribute to enzyme interactions.

### 4. Resistance to auranofin is not conferred by an isolated mutation in TgSOD2

The anti-parasitic effect of auranofin appears to stem from its molecule of gold, which is known to induce the accumulation of ROS in parasites (4, 34). Since we hypothesized that a single mutation (SNV) suffices to generate resistance to auranofin, we analyzed the effect of the TgSOD2 L201P mutation on auranofin sensitivity. SOD2 is a *bona fide* enzymatic ROS scavenger and SOD2 is the sole mitochondrial superoxide dismutase in tachyzoites. To analyze the role of mutant SOD2 in auranofin resistance, we generated a *T. gondii* clone with a knock-in YFP tag for SOD2, that localizes to the tachyzoite mitochondrion as previously reported (Figure 5A). Subsequently, we generated two *T. gondii* SOD2.L201P lines: one lacking the YFP tag and one tagged with a C-terminal YFP fusion (Figure S1*/* Both lines were corroborated by Sanger sequencing. In the absence of Auranofin, both L201P lines grew less well than the TgSOD2-YFP wild-type line but were more fit than Aur^R^ line 8 (Figure 5B). Previous attempts to create SOD1 and SOD2 gene deletions failed (35) and SOD2 is a fitness-conferring locus in a genome-wide CRISPR/Cas screen (36). It is noteworthy that the L201P mutation is associated with reduced fitness in the absence of superoxide stress (control conditions). In the presence of auranofin, only line 8 grows well, indicating that the single L201P mutation to TgSOD2 does not confer resistance. This may indicate that this substitution does not influence superoxide metabolism at all or that auranofin resistance also requires the mutant TgRNR enzyme. It is unlikely that these enzymes directly interact, as RNR enzymes are cytosolic while TgSOD2 is in the mitochondrial compartment. However, the SOD2.L201P *T. gondii* line accumulated ROS similarly to wild-type parasites, consistent with its auranofin-sensitive growth (Figure 5C).

**Figure 5:**
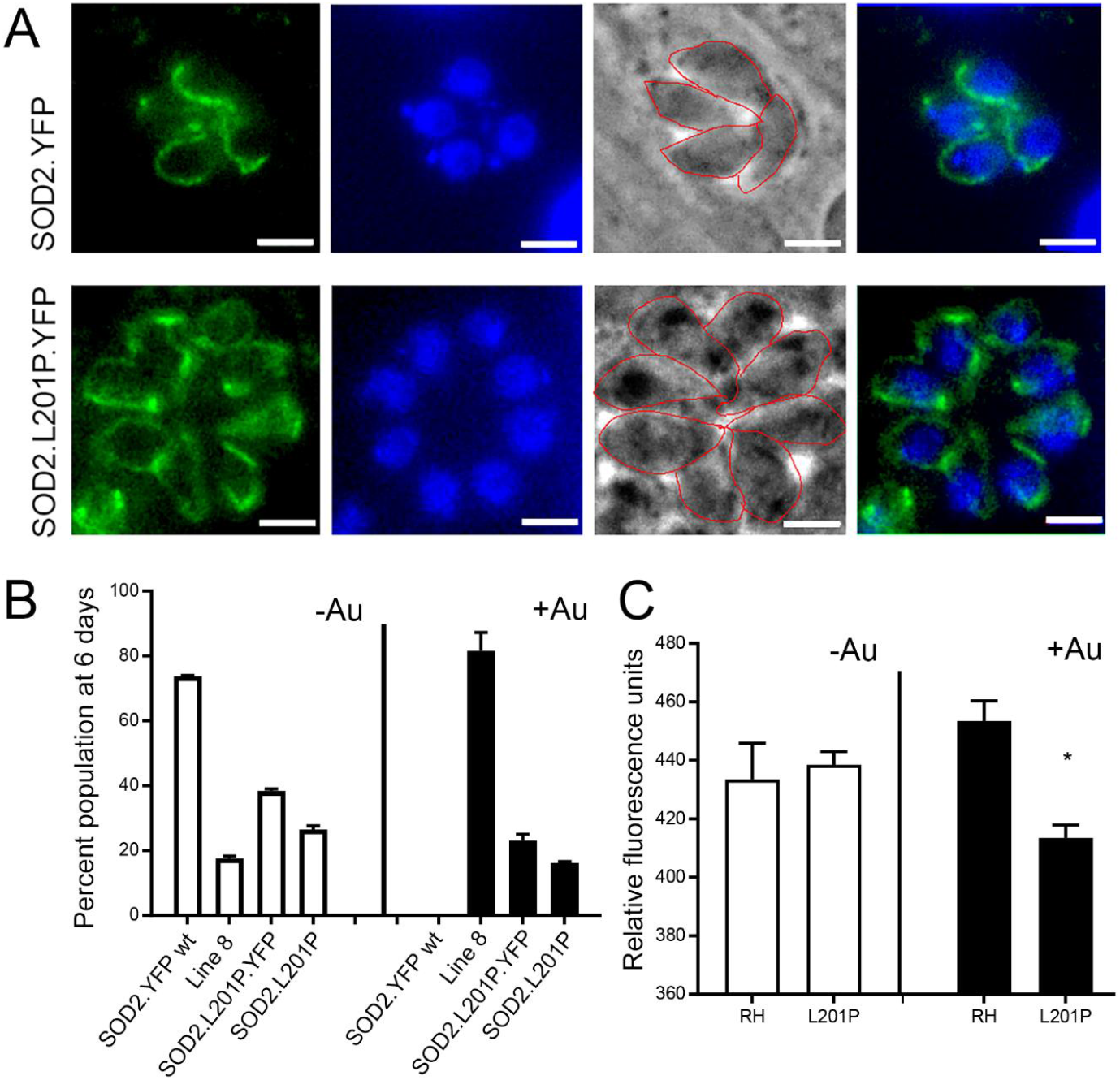
A L201P mutation in TgSOD2 does not alter localization of TgSOD2 or confer resistance to auranofin. (A) Wild type and mutant SOD2-YFP exhibit a mitochondrial distribution. (B) Growth competition assays indicate that the L201P mutation is associated with reduced fitness in the absence of auranofin (control conditions, white bars). In the presence of auranofin (black bars), only line 8 shows growth advantage, indicating that the single L201P mutation to TgSOD2 does not confer resistance. (C) In the absence of auranofin, the SOD2.L201P line accumulated ROS similarly to wild type parasites, consistent with its auranofin-sensitive growth; however, it accumulated less ROS in the presence of auranofin

## Discussion

Auranofin, is an FDA-approved drug for treatment of rheumatoid arthritis. More recently it has been proposed to be a promising anti-parasitic agent with activity against diverse parasites with public health significance. We previously demonstrated that auranofin decreases the parasite burden during a systemic infection with *T. gondii* using a chicken embryo model of acute toxoplasmosis (13). Our preliminary observations suggest that auranofin exerts a dual effect: it modulates both parasite proliferation and host immunopathology. Thioredoxin reductase has been implicated to be the molecular target of auranofin in parasites. Herein, we report isolation and characterization of auranofin-resistant *T. gondii* lines. None of our resistant lines has mutations in the thioredoxin reductase gene. This may indicate that this enzyme is not the molecular target of auranofin in *T. gondii;* alternately, it may indicate that thioredoxin reductase cannot tolerate resistance-conferring mutations or is not the sole significant target in this organism. Each of our auranofin resistant *T. gondii* lines carry several mutations (SNVs). Importantly, resistant lines accumulate less reactive oxygen species (ROS) than wild-type parasites.

There are several approaches to target identification in drug discovery. In *T. gondii*, chemical mutagenesis is a well-established and robust approach (*i.e*. (15, 19) to identify the molecular targets of candidate anti-*T. gondii* drugs. This approach is also relevant to anticipating mechanisms of drug resistance. We isolated auranofin-resistant *T. gondii* lines derived from the RH strain. Resistant lines have increased replication relative to wild type RH parasites in competition assays carried out in the presence of auranofin. Lines with ≤5 accumulated SNVs (lines 1-4) appear to have a growth advantage over wild-type parasites in the absence auranofin. This probably reflects a fitness defect induced by expression of YFP in wild type parasites (21); these lines likely grow comparably to non-YFP expressing wild type parasites without auranofin present. In contrast, resistant lines that have >5 SNVs each compete poorly with wild type parasites in the absence of selection, suggesting that accumulation of SNVs comes at a cost to parasite fitness. An alternative explanation to the isolation of multiple SNVs in each resistant clone is that auranofin interferes with many processes and development of resistance requires alterations to more than one target.

Auranofin contains a molecule of gold in a 3,4,5-Triacetyloxy-6-sulfanyl-oxan-2-yl methyl ethanoate scaffold and has been proposed to be a pro-drug (37) that delivers gold to dithiol groups, like those found in proteins that bear thioredoxin-containing domains (38). Although auranofin inhibits mammalian thioredoxin reductase (39), the source of its anti-inflammatory activity remains ill-defined and may reflect inhibition of multiple targets. For example, macrophages decrease production of nitric oxide and pro-inflammatory cytokines in response to auranofin, which also inhibits cyclooxygenase-2 -dependent prostaglandin E2 production (40). Previous studies using the parasite enzymes trypanothione reductase and glutathione-thioredoxin reductase incubated with auranofin demonstrated that co-crystals harbor gold bound to reactive cysteines in these proteins (2, 3, 41) In addition, *S. mansoni* worms with decreased expression of thioredoxin glutathione reductase (SmTGR) exhibit diminished parasite viability upon exposure to auranofin. *E. histolytica* treated with auranofin upregulate transcripts of a gene encoding a protein resembling an arsenite-inducible RNA-associated protein (AIRAP) (4). Interestingly, both arsenite and auranofin are inhibitors of thioredoxin reductase, implying that EhTrxR is a target of auranofin. Studies in both *S. mansoni* and *E. histolytica* corroborated the interaction of thioredoxin reductase with auranofin by mobility shift assays or other in vitro measures (4, 14). In these cases, the approach involved target validation guided by the chemical properties of gold rather than target identification through genetic resistance.

It was surprising to not identify changes to the thioredoxin reductase gene, given previous studies that implicate antioxidant enzymes containing thioredoxin or thioredoxin-like domains as a target for auranofin in parasites. However, although *S. mansoni* and *E. histolytica* thioredoxin reductase antioxidant systems are essential for survival, a previous study indicates that *T. gondii* thioredoxin reductase is not essential (42). It should be noted that infection with 10^6^ TgTrxR knockout parasites increases survival in mice by approximately 2 weeks relative to the parental RH line (42). *In vitro*, the null line exhibits reduced antioxidant capacity, invasion efficiency, and proliferation. These observations are in-line with the CRISPR fitness score for thioredoxin reductase (−1.98) which places it as fitness-conferring but not essential.

Among the significant SNVs found in our auranofin resistant *T. gondii* clones, we directed our attention at a point mutation to the superoxide dismutase 2 (SOD2) gene. SOD2 is a *bona fide* enzymatic antioxidant system that catalyzes the dismutation of superoxide anions into hydrogen peroxide and oxygen. The *Toxoplasma* genome contains three SOD genes. TgSOD1 is expressed in the cytoplasm of tachyzoites and TgSOD2 is located in the mitochondrion; TgSOD3 is specifically expressed in oocysts. TgSOD2 is a critical enzyme for tachyzoite survival: previous efforts to generate a TgSOD2 knockout line were not successful and its CRISPR fitness score for (−4.09) indicates that it is strongly fitness conferring and likely essential (35). We hypothesized that the L201P mutation in TgSOD2 might increase its capacity to convert superoxide into hydrogen peroxide and oxygen to confer auranofin resistance. To test this hypothesis, we generated several lines that harbor knock-in tag to wild type SOD2 and engineered both YFP-tagged and untagged lines that bear the L201P mutation. Consistent with previous characterization of TgSOD2 (32), both wild type and L201P TgSOD2-YFP lines exhibit a mitochondrial distribution. However, neither the tagged or untagged L201P TgSOD2 lines exhibit increased growth in 2.5 uM auranofin, indicating that the single TgSOD2 mutation cannot confer resistance to this level of auranofin or reduce accumulation of ROS in our assay.

In addition to the L201P TgSOD2 mutation, Aur^R^ line 8 harbors a mutation in the large subunit of TgRNR, an essential enzyme which generates deoxyribonucleotides that are required for genome duplication and parasite replication. TgRNR has a 59 amino acid N-terminal extension relative to other RNRs. Peptides from this extension are represented in phosphorylated and monomethylated MS datasets, suggesting that this domain may regulate RNR activity. RNRs perform free-radical chemistry, require electrons donated from thioredoxin and are susceptible to superoxide inactivation. Although the L166Q mutation is in an area that is not known to contribute to enzyme interactions, it may play a role diminishing TgRNR susceptibility to superoxide inactivation. While this mutation may indirectly contribute to resistance to superoxide, RNR enzymes do not directly contribute to redox homeostasis and the observed reduced accumulation of ROS in the Aur^R^ lines.

Our previous and current findings suggest that auranofin decreases *T. gondii* viability through oxidative stress (13). Aur^R^ lines exhibit an increased ability to scavenge ROS. The genetic studies presented here did not identify a consistent mechanism for acquisition of resistance by point mutations to a specific protein target. Our future studies will examine whether there are transcriptional, translational or post-translational changes that contribute to auranofin resistance in these lines. For example, difference-gel electrophoresis and mass spectrometry have been used to identify differentially expressed proteins in sulfadiazine-resistant *T. gondii* isolates. While it was disappointing to not identify a single molecular target for auranofin in *T. gondii* in our studies, this result likely indicates that an enhanced ability to scavenge ROS in the presence of auranofin requires multiple changes to the parasite genome. Therefore, if auranofin is repurposed as an anti-parasitic treatment, it would likely present an increased threshold to resistance in clinical contexts.

## SUPPLEMENTAL FIGURES

**Figure S1:**
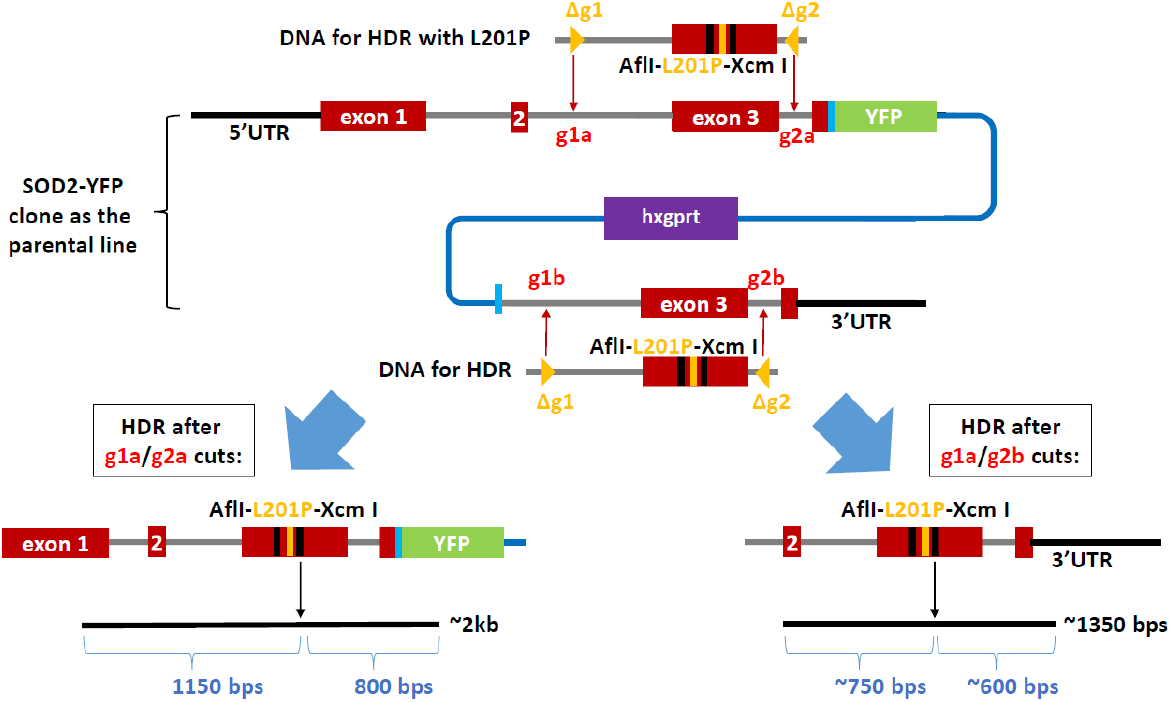
CRISPR-Cas9 strategy for generating the *T. gondii* SOD2.L201P line. TgSOD2-YFP parasites incorporating the LIC plasmid (blue line) undergo CRISPR-Cas9 mediated homologous directed repair (HDR). The DNA amplicon used for HDR contains the L201P mutation (gold vertical line in exon 3) and two silent and unique restriction sites AflI and XcmI (black vertical lines in exon 3), which allow for screening of positive clones. Sequences corresponding to the two gRNA were deleted from the DNA for HDR (Δg1 and Δg2, golden triangles). Each of the gRNA was present twice in TgSDO2-YFP parasites (g1a/b, g2a/b; red font), resulting in two *T. gondii* SOD2.L201P lines. For HDR after g1a/g2a cuts, a *T. gondii* SOD2.YFP.L201P line is generated (lower left). For HDR after g1a/g2b cuts, a *T. gondii* SOD2.L201P line without YFP or LIC plasmid (blue) is generated (lower right). Positive clones were screened with XcmI digest before sequence verification.

**Table S1:**
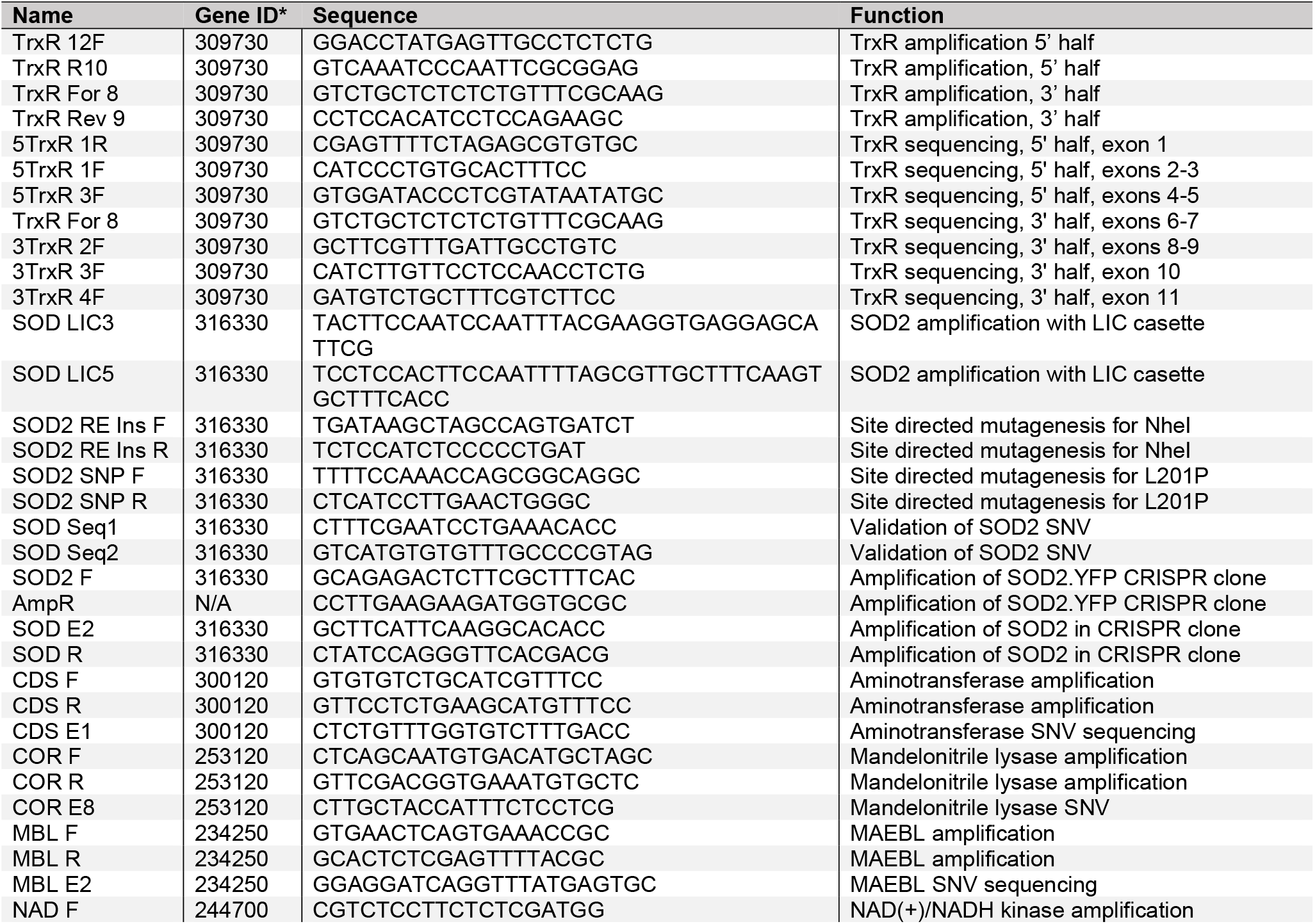
Primers for SNV validation

## Acknowledgements

Special gratitude to Dr. Melissa Lodoen (UCI) for her advice and critical review of this manuscript and to Edward Brignole (MIT) for his thoughts on the TgRNR mutation. RMA was supported by NIHK08 5K08AI102989-04 and AMFDP RWJF 70642 (the views expressed here do not necessarily reflect the views of the Foundation). This work was made possible, in part, through access to the Genomics High Throughput Facility Shared Resource of the Cancer Center Support Grant (P30CA-062203) at the University of California, Irvine and NIH shared instrumentation grants 1S10RR025496-01, 1S10OD010794-01, and 1S10OD021718-01.

